# Taking advantage of external mechanical work to reduce metabolic cost: the mechanics and energetics of split-belt treadmill walking

**DOI:** 10.1101/500835

**Authors:** Natalia Sánchez, Surabhi N. Simha, J. Maxwell Donelan, James M. Finley

**Affiliations:** Division of Biokinesiology and Physical Therapy, University of Southern California, Los Angeles, CA, 90033; Department of Biomedical Physiology and Kinesiology, Simon Fraser University, Burnaby, British Columbia, Canada V5A 1S6; Department of Biomedical Engineering, University of Southern California, Los Angeles, CA, 90089; Neuroscience Graduate Program, University of Southern California, Los Angeles, CA, 90089

**Author notes:** Denotes equal contributions (Natalia Sánchez, Surabhi N. Simha). **Author Email Addresses:** Natalia Sanchez, Surabhi Simha, J. Maxwell Donelan, James M. Finley. **Mailing Address:** University of Southern California, Division of Biokinesiology and Physical Therapy 1540 E. Alcazar St, CHP 155 Los Angeles, CA, USA 90033.

## Abstract

In everyday tasks such as walking and running, we exploit the work performed by external sources such as gravity to reduce the work performed by muscles. There has been considerable recent effort to design devices capable of performing mechanical work to improve walking function or reduce effort. The success of these devices relies on the user adapting their natural control strategies to take advantage of assistance provided by the device. Although locomotor adaptation is central to this process, the study of adaptation is often done using approaches that on the surface, seem to have little in common with the use of external assistance. Here, we show that one of the most common approaches for studying this process, which is adaptation to walking on a split-belt treadmill, can be understood from a perspective in which people learn to take advantage of mechanical work performed by the treadmill. During adaptation, people systematically adjust their step lengths, defined as the distance between the feet at heel strike, from one step to the next. Initially, the step length on the slow belt is longer than the step length on the fast belt, measured as a negative step length asymmetry, but people naturally reduce this asymmetry with practice. Here, we demonstrate that these modifications of step length asymmetry allow people to extract positive work from the treadmill belts to reduce the positive work performed by the legs and simultaneously reduce metabolic cost. Moreover, we show that walking with a positive step length asymmetry minimizes metabolic cost, and people prefer to walk in this manner when allowed to select their walking pattern. Together, our results suggest that split-belt adaptation can be interpreted as a process by which people learn to take advantage of mechanical work performed by an external device to improve walking economy.

## Introduction

The nervous system can learn to take advantage of external assistance to produce and sustain motion. In everyday tasks such as walking and running, we often exploit the passive dynamics that arise from the interaction between the body and gravity to reduce the need for muscle force [1,2]. External assistance of our motion is not restricted to gravity—in surfing we learn to harvest the assistance of waves and in sailing we learn to harvest the assistance of the wind. The ability of our nervous system to harness energy is critical if we are to take advantage of assistance from devices like powered prostheses and exoskeletons. Exoskeletons for the lower limbs are often designed to reduce muscular work, reduce effort, and increase endurance during walking [3–5]. These devices commonly use powered actuators [6–13], designed to reduce metabolic cost by performing mechanical work that would otherwise need to be generated by muscles. However, the degree to which these assistive devices reduce metabolic cost depends not only on the amount of work performed by the device, but also on the individual’s ability to exploit adaptive learning processes to take advantage of the external assistance [14].

Although learning is key to maximizing the benefits from external assistance, the study of locomotor learning is often done in contexts that do not seem to have much in common with the study of assistive devices. For example, much of the work on adaptive locomotor learning uses a split-belt treadmill paradigm where individuals walk on a treadmill with two belts that move at different speeds [15–17]. Upon initial exposure to walking with the belts moving at different speeds, the distance between the feet at leading limb heel-strike—referred to as step length—becomes asymmetric. The step length at slow leg heel-strike becomes longer than the step length at the fast leg heel-strike because the fast belt pulls the leg into a more extended position. This asymmetry in step lengths is gradually reduced over the course of 10 to 15 minutes [15] and is accompanied by a reduction in positive mechanical work performed by the legs [18] and a reduction in metabolic cost [19]. A potential explanation for these observations is that individuals may adapt their gait in split-belt walking to minimize metabolic cost [19,20].

Here we use principles from mechanics to illustrate that, like exoskeletons, split-belt treadmills can provide assistance during walking. As we detail in the Theory and Predictions section, gaining assistance in the form of net mechanical work on the person from the treadmill is unique to the split-belt treadmill and is not possible on a normal “tied-belt” treadmill, or when walking over ground. Walking at a constant speed in any of these situations requires that the person generates braking and propulsive impulses that are balanced throughout the gait cycle. On a split belt treadmill, however, people can choose how to distribute braking and propulsion between the two belts to take advantage of the difference in belt speeds. If the forces and work generated by the legs are redistributed properly, the work performed by the treadmill on the person could be used to reduce the positive work required by the person’s muscles and ultimately reduce metabolic cost.

Split-belt treadmills provide a unique approach to study how to gain advantage of external assistance, as individuals could reduce the energetic cost of walking by stepping further forward on to the fast belt relative to the slow belt [21]. As we describe in Theory and Predictions, this pattern would generate a larger braking force on the fast belt, which would be balanced by more propulsive force applied to the slow belt. Because of the difference in belt speeds, this results in net negative work performed by the person on the belts, and net positive work performed by the belts on the person. It is not guaranteed that the person benefits from this positive treadmill work—they may dissipate it by performing additional negative work. But it is also possible that they allow the positive work from the belts to assist walking by increasing their mechanical energy, reducing the positive mechanical work required from their muscles, and reducing their metabolic cost. This can only occur if individuals learn to take advantage of the treadmill work by reducing positive muscle work. Here, we show that individuals can learn to adapt their coordination to take advantage of the resulting positive work performed by the treadmill and reduce metabolic cost.

### Theory and Predictions

Here we consider three walking conditions: ground walking, walking on side-by-side treadmills with belts “tied” to move at the same speed, and walking on side-by-side treadmills with belts “split” to move at different speeds. In ground walking, the ground is level and the person walks at a constant average speed relative to the ground. In the two treadmill cases, the treadmills are level and the person keeps the same average position on the treadmills equating to zero average speed relative to the stationary ground. (It is perhaps easiest to understand the following arguments when considered from a reference frame that is fixed to the ground, and not to one of the treadmill belts. The arguments do hold for other reference frames, as long as they are not accelerating and as long as one does not switch between reference frames.) In the treadmill conditions, we consider three systems: the person, the left treadmill, and the right treadmill. The left and right treadmills move at the same speed in the tied-belt condition, and at different speeds in the split-belt condition. In the ground walking condition, the ground has zero speed resulting in only two systems: the person and the ground. In all of the above walking conditions, the following two constraints must be fulfilled.

C1. The sum of the external forces acting on the person must be zero on average. Otherwise, there would be net acceleration or net vertical displacement violating the requirements of steady-state speed and level walking.
C2. The net mechanical work on the person must be zero on average. Otherwise, there would be a net gain in kinetic or potential energy, again violating the steady-state speed and level walking requirements. Importantly, these two constraints taken together do not mean that external forces (e.g. treadmill forces) cannot perform net mechanical work on the person. Indeed, they could perform net negative or net positive work while still summing to zero net system acceleration as long as forces internal to the person system (e.g. muscle forces) perform the opposite amount of work.

These constraints affect the three walking conditions in different ways. For ground walking, they mean that the work performed by the person must be zero on average. This is because the ground cannot perform work on the person—relative to the ground-fixed reference frame, there is no displacement at the ground point of force application.

Neglecting air resistance, the only source of work is the person and thus the person must perform no net work (C2). Performing no net work can be accomplished by performing zero work, but more likely it is accomplished with equal amounts of positive and negative work. To assist with understanding, consider the person as a point mass body with legs that are massless pistons that generate forces on the ground (or treadmill belts as in Figure 1A & B) and equal but opposite forces on the point mass. In this model, all the work performed by the person is performed by the legs. In ground walking, the legs can perform both positive and negative work as long as they sum to zero.

**Figure 1.**
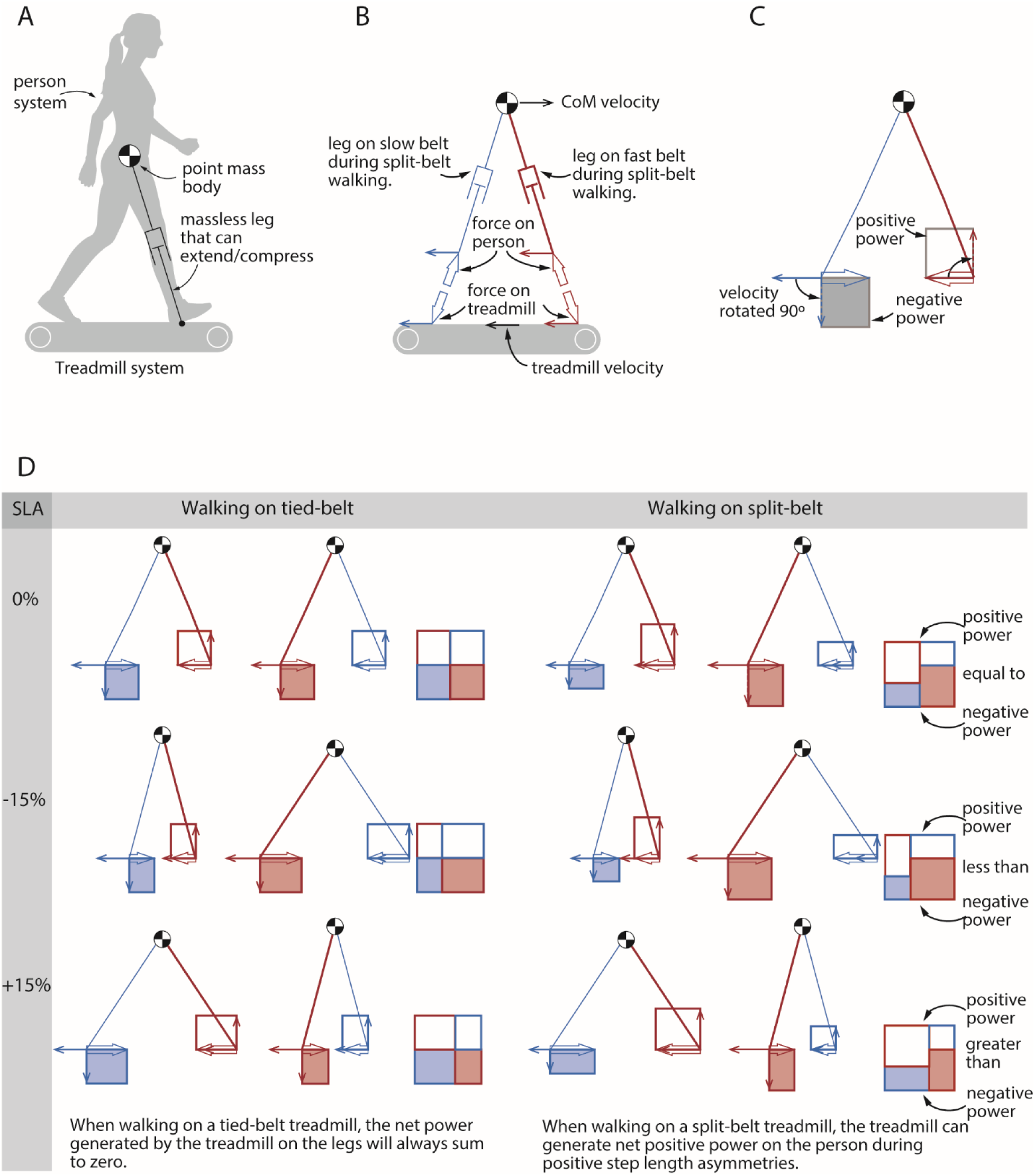
A. We model the person as a point mass body with massless legs that can extend or compress. B. Free-body diagram of the person and treadmill systems. Blue represents the leg on the slow belt and red represents the leg on the fast belt when the belt velocities are split. Large open arrows represent forces, line arrows represent velocities. C. Power generated by the treadmill on the legs. Dashed arrows represent velocities rotated by 90 degrees. We use the individual limbs method to calculate the power generated by the treadmill on each leg during a stride (not shown). The positive power generated by the treadmill on the legs is visualized with open rectangles above the point of force application, and negative power with closed rectangles below the point of force application. D. Power generated by the treadmill on the legs. During tied belt walking, the positive power generated by the treadmill on the legs is always equal to the negative power generated within a stride, at all step length asymmetries (SLA). This is true even though the total power generated during the long step is greater than that generated during the short step, at asymmetric step lengths. We can observe this through the rectangles shown to the right of each condition, where the sum of the top open rectangles always equals the sum of the bottom closed rectangles. However, when walking on split-belt treadmill, we see that at positive step length asymmetries (bottom row), the treadmill generates net positive power on the legs during the long step that is greater than the net negative power it generates during the short step. This leads to a net positive power on the person over the stride.

The constraints explained above also require the person to perform zero net work during tied-belt treadmill walking. Unlike ground walking, the person performs work on the belts during treadmill walking when considered from a ground-fixed reference frame. At heel contact, for example, the force exerted by the leading leg opposes the belt velocity at the point of contact (Figure 1B). Since the velocity at the point of force application only has a fore-aft component, the power generated by the treadmill on the leg is the dot product of the horizontal fore-aft force from the treadmill and the treadmill velocity. It is positive when the force and velocity are in the same direction, and negative when the two are opposite. At heel contact, this force will perform negative work on the belt. The belt does not slow down because its motor simultaneously does an equal amount of positive work on the belt. (For the sake of building intuition, we assume an ideal treadmill where powerful motors are under rapid feedback control to keep belt speed constant irrespective of the forces that the legs apply to it.) The reaction force of the belt on the person is equal and opposite to the force of the person on the belt (Figure 1B), but the velocity of the point of force application is the same—when a person does negative work on a belt, the belt does an equal amount of positive work on the person (Figure 1C).

When the belts are moving at the same speed, as is the case for the tied-belt condition, the positive and negative work done by the person on the belts, and that done by the belts on the person, both must sum to zero. This is because of the constraint that the external forces must sum to be zero on average (C1). In the fore-aft direction, for example, if at some point in the stride the person generates a force on a belt that has a braking component, the person must generate an equal propulsive force at some other point during the stride to prevent drift on the treadmill. Since the belt speeds are equal, balancing forces also means balancing the work done by the person on the belts, and by the belts on the person. That is, a person’s braking force will get the benefit of positive work being done on the person by the belts that will be counteracted when the person generates an equal propulsive force resulting in the belts doing negative work on the person. Thus, a person walking on tied belts cannot benefit from net work performed by the belts on the person—it will always sum to zero (Figure 1D). Another way to make this point is that our choice of reference frame is arbitrary as long as it is not accelerating. Thus, treadmill walking in a belt-fixed reference frame is indistinguishable to over-ground walking in a ground-fixed reference frame.

In contrast to ground and tied-belt walking, in split-belt walking the belts can do net work on the person, and the work done by the person does not need to sum to zero. The external forces acting on the person must still average zero (C1), and the net work on the person must also average zero (C2). This zero net work may be accomplished with net positive or negative work by the person and net negative or positive work by the treadmills (Figure 1D). For example, the person’s fast leg might provide net propulsive force on the fast belt resulting in net positive fast leg work, and then the person’s slow leg might provide an equal amount of net braking force on the slow belt to bring the average horizontal force to zero. However, because of the differences in belt speeds, this braking force would require the slow leg to perform less net negative work on the slow belt than the net positive work that the fast leg had to perform on the fast belt for the same magnitude horizontal force. Together, the person’s legs would perform net positive work on the belts, and consequently, the belts would perform net negative work on the person. Alternatively, if the fast leg was responsible for more of the braking, and the slow leg for more of the propulsion, the person would perform net negative work and the person would gain net positive work from the treadmill belts.

Based on these predictions, a split-belt treadmill can be viewed as an assistive device— similar to an exoskeleton—where the person has to learn how to coordinate their legs to maximize the assistance from the motors. Positive muscle mechanical work is metabolically expensive relative to negative muscle mechanical work requiring roughly five times the ATP per Joule [22,23]. To decrease the relatively expensive positive mechanical work, a person should decrease the propulsive forces generated by the fast leg on the fast belt, and instead rely on the slow leg to perform more propulsion for less positive work. Assuming the legs generate forces that act approximately along the legs, the person can reduce fast leg propulsive forces by lifting off the fast belt with less hip extension or contacting during heel strike at a greater angle with respect to the vertical, and the person can increase slow leg propulsive forces by lifting off at a larger hip extension angle and contacting at a smaller angle during heel strike. All else being equal, this effect will shorten the distance between the feet at slow-leg heel-strike (slow step length: *SL*_*Slow*_) and lengthen the distance between the feet at fast-leg heel-strike (fast step length *SL*_*Fast*_.) If we consider step length asymmetry as a measure of the difference between the fast step length and the slow step length, this change in coordination will shift asymmetry to more positive values.

To summarize, we predict that as an individual adopts a more positive step length asymmetry, the split-belt treadmill will perform more positive work on the person, reducing the positive work required by their muscles. Because of the relative expense of positive work, we also predict a reduction in the metabolic cost of walking as step length asymmetry becomes more positive, consistent with the findings from previous adaptation experiments [19]. However, we do not attempt to predict the metabolically optimal step length asymmetry as positive muscle mechanical work is only one contributor to metabolic cost, and it is not clear how other contributors, such as the cost to swing the legs [24], or the cost of maintaining an upright posture [25], change with step length asymmetry.

## Results

We determined whether individuals can modify their step lengths while walking on a split-belt treadmill to gain assistance from the treadmill, reduce positive leg work, and ultimately reduce metabolic cost. To this end, we mapped the relationship between step length asymmetry, mechanical work, and metabolic cost in 16 healthy participants while walking on a dual belt instrumented treadmill. The left and right belt speeds were set at 1.5 m/s and 0.5 m/s, respectively, and participants used visual feedback to maintain step length asymmetries of 0.00, +/-0.05, +/-0.10 and +/-0.15 in separate, six-minute trials presented in a random order (Fig. 2A-B). Positive asymmetries indicate longer steps on the fast belt, while negative asymmetries indicate longer steps on the slow belt. We computed the mechanical work performed by the legs and the treadmill using an extension of the individual limbs method [18,26] and we calculated metabolic rate from expired gas analysis using the Brockway equation [27]. Lastly, we used mixed-effect regression models to determine 1) if the positive work performed by the treadmill increased as step length asymmetry became more positive, 2) if people take advantage of the positive work performed by the treadmill to reduce the positive work performed by the legs, and 3) whether reductions in positive work performed by the legs are accompanied by reductions in metabolic cost.

**Figure 1:**
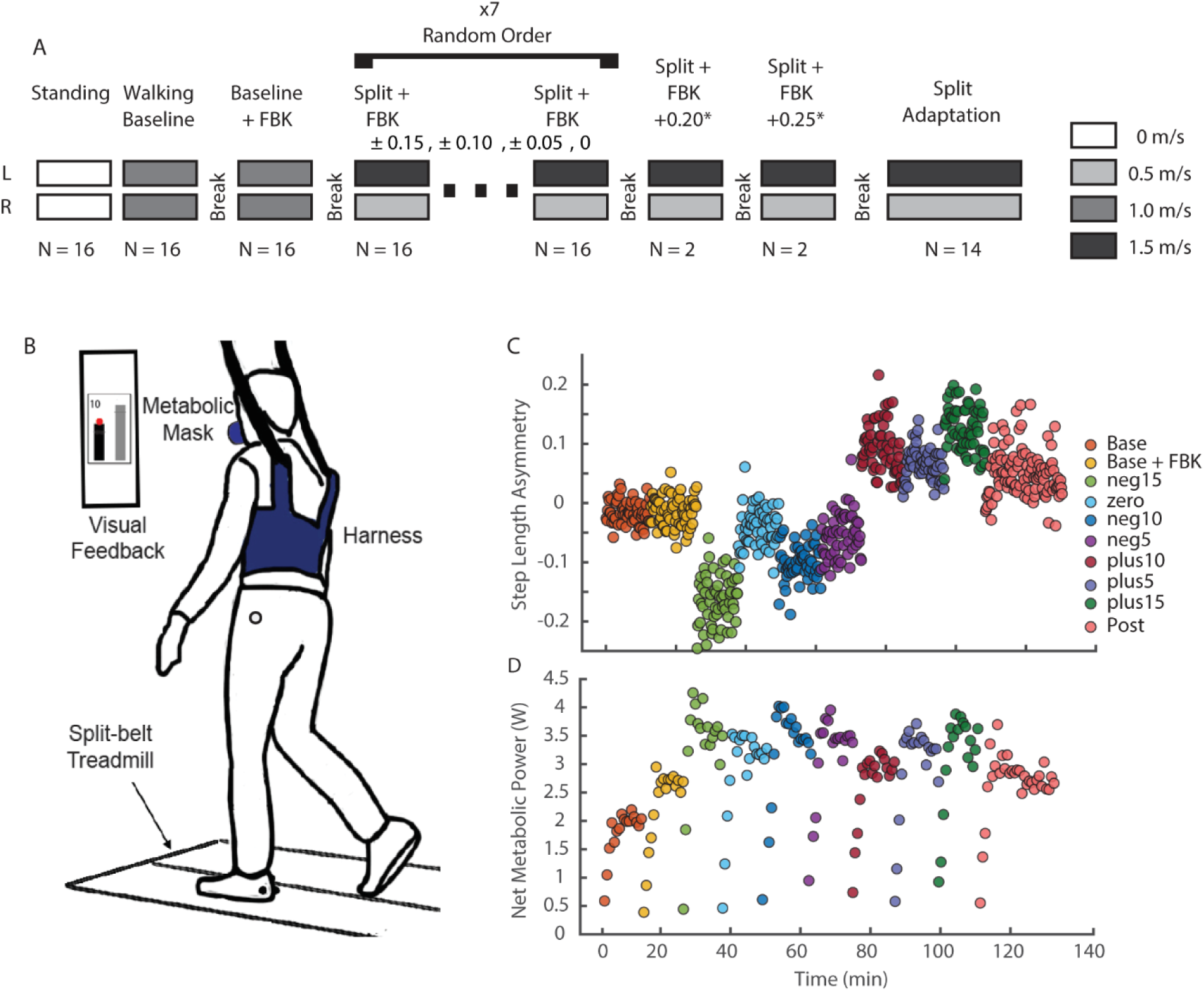
Experimental protocol and setup. A) Detailed experimental protocol. We randomized the order of the split-belt trials during which feedback of step lengths was provided to obtain seven different levels of asymmetry. Two participants performed two additional trials for levels of 0.20 and 0.25 step length asymmetry. All participants performed the split-belt adaptation trial last. The adaptation trial lasted 10 minutes; all other trials lasted six minutes with four-minute breaks in between. B) Experimental setup. Reflective markers were placed bilaterally on the lateral malleoli to measure step lengths in real time. Two additional markers were placed on left and right greater trochanters to measure hip width for the visual feedback. An overhead harness was used to prevent falls without providing body weight support. No handrail was provided. Metabolic cost was measured using expired gas analyses. C) Raw values of step length asymmetry for all walking trials for a representative participant. Average step length asymmetry was binned every 10s to illustrate performance in the time domain. D) Raw net metabolic power values measured for each walking trial for the participant in panel C. All values were baseline corrected to standing baseline.

### Modulation of Foot Position to Achieve Target Step Length Asymmetry

Participants modified the configuration of both the leading and trailing limbs as they varied step length asymmetry throughout the experiment. We measured changes in limb configuration by computing the distance between the feet and an estimate of the body’s center of mass. During the visual feedback conditions, as asymmetry increased towards positive values, participants lengthened the step on the fast belt by stepping further forward in front of the center of mass (Figure 3A, β = 243.4, 95% CI [212.2, 274.6], p=2.22e-29) and allowing the slow leg to trail further behind the center of mass (Figure 3A, β = -356.1, 95% CI [-385.1, -327.1], p=4.43e-46). As asymmetry became more positive, participants shortened the slow step by placing the slow leg closer to the center of mass (Figure 3B, β = -348.5, 95% CI [-385.3, -311.6], p=5.00e-36) and making minor adjustments in the position of the trailing limb (Figure 3B, β = 85.4, 95% CI [56.5, 114.3], p=5.17e-8).

**Figure 3:**
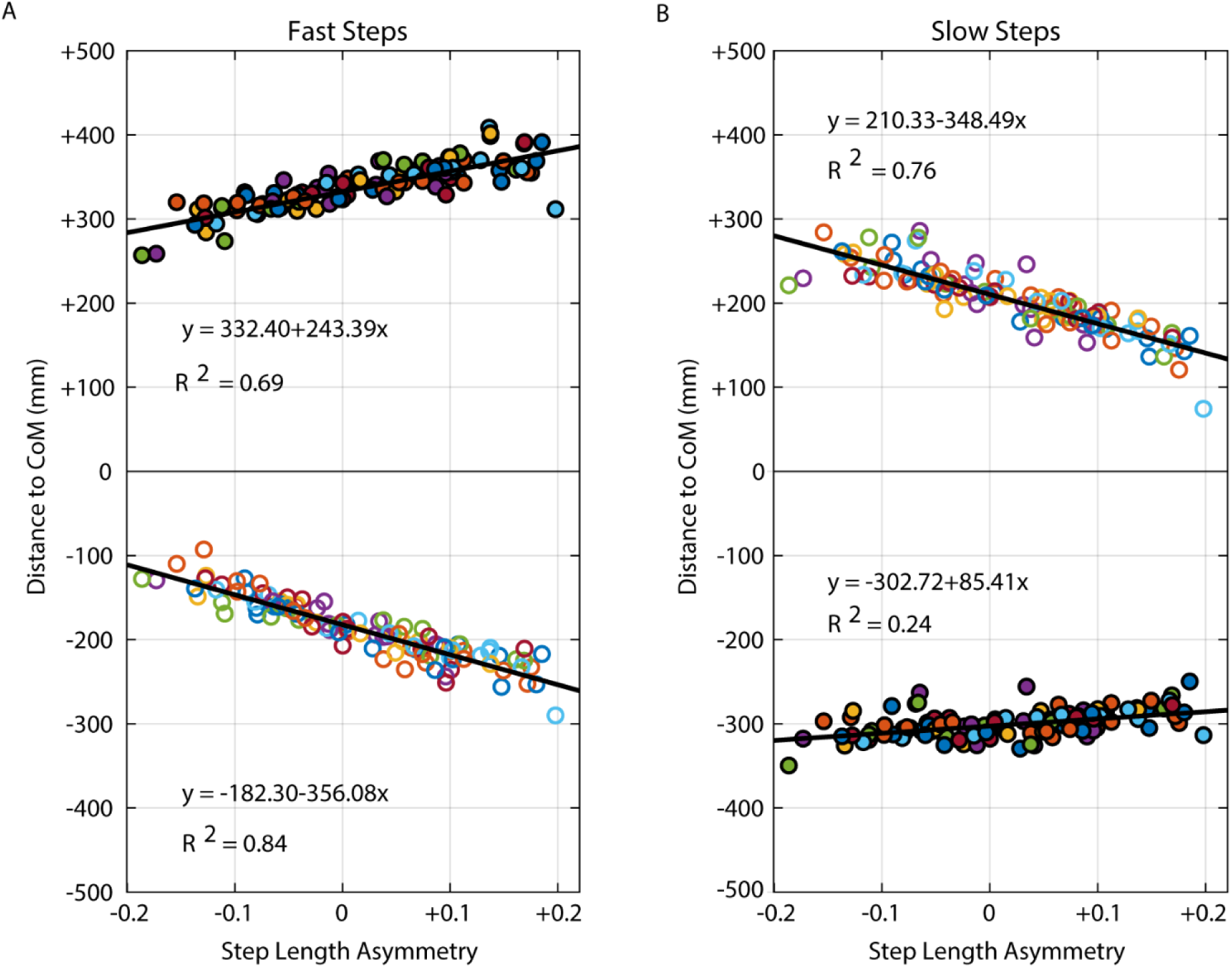
Distance from the leading and trailing foot to the center of mass at heel strike. Positive values indicate that the foot is anterior to the center of mass while negative values indicate that the foot is posterior to the center of mass. Closed and open points correspond to the fast and slow limb, respectively. The distance between the leading and trailing feet at heel strike constitutes the step length. A) Fast step lengths. To increase the fast step length as asymmetry increased from negative to positive, participants increased both leading foot distance to the center of mass and trailing foot distance to the center of mass. B) Slow step lengths. To decrease the slow step length as asymmetry increased from negative to positive, participants primarily decreased the distance from the leading foot to the center of mass while maintaining a relatively consistent trailing foot position.

### Mechanical Work Performed by the Legs and the Treadmill Varies with Step Length Asymmetry

Consistent with our hypothesis, participants transitioned from performing net positive work with the legs at negative step length asymmetries to performing net negative work at positive asymmetries. This change in work performed by the limbs can be appreciated by understanding how the mechanical power generated by each limb changes across levels of asymmetry. The fast leg transitioned from generating a large amount of positive power during push-off at -15% asymmetry to performing a large amount of negative work during weight acceptance at an asymmetry of +15% (Figure 4A). In contrast, there were negligible changes in mechanical power in the slow leg across levels of asymmetry (Figure 4B). Overall, the amount of positive work performed by the legs decreased by ∼13% between - 15% and +15% step length asymmetry (Figure 5A, β = -0.009, 95% CI [-0.013, -0.006], p=3.52e-7). In contrast, the amount of negative work performed by the legs increased by ∼33% over the full range of step length asymmetries (Figure 5B, β = -0.019, 95% CI [-0.0220, -0.0151], p=7.59e-19). As a result of these combined changes, the legs ultimately performed more negative work than positive work at positive asymmetries (Figure 5C, β = -0.028, 95% CI [-0.029, -0.027], p=1.66e-67).

**Figure 4:**
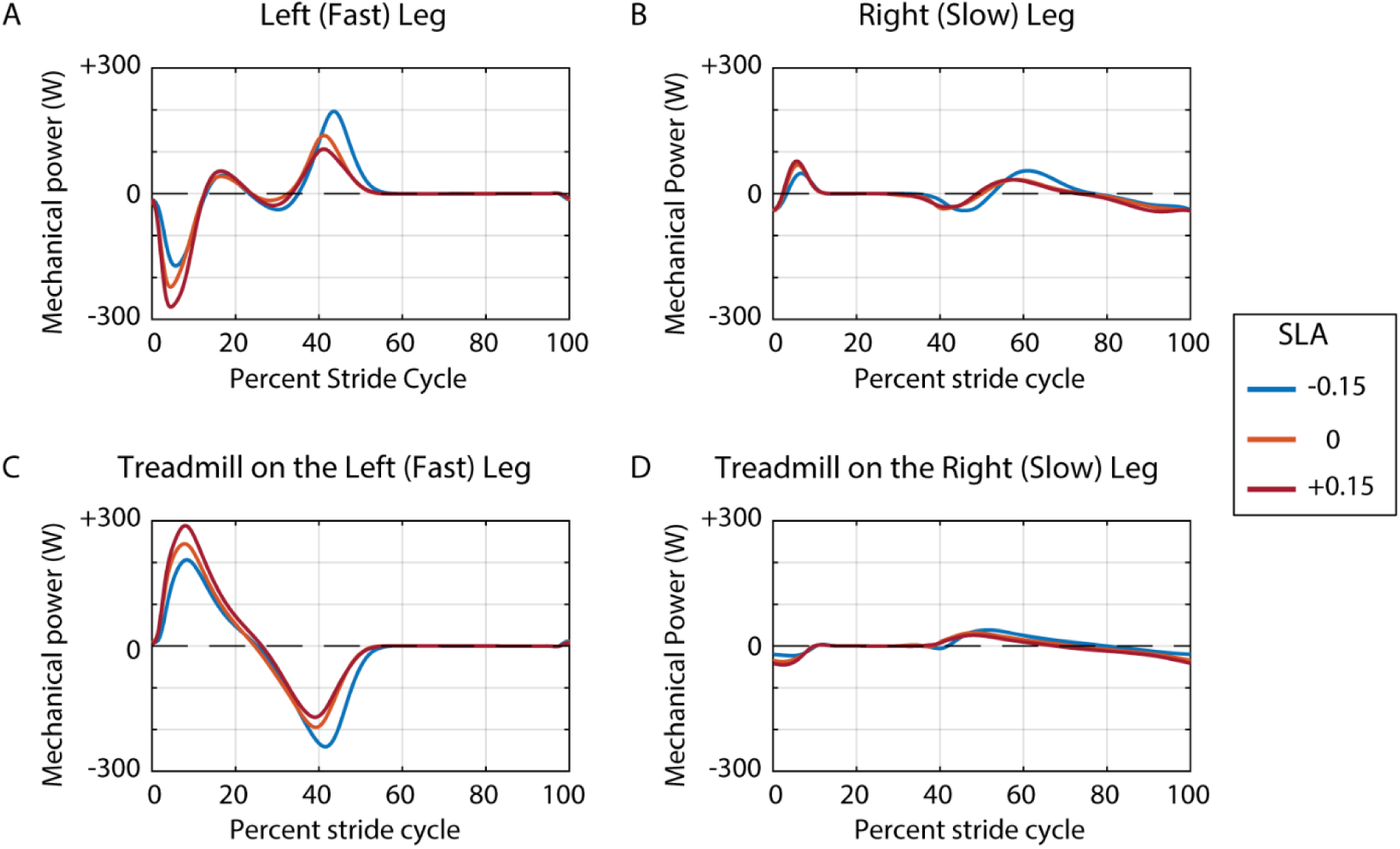
Mechanical power generated by the A) fast leg, B) slow leg, C) fast belt, and D) slow belt throughout the stride cycle. The power generated by the treadmill belts represents the rate of mechanical work performed by each belt on the body. During the early portion of the stride cycle (∼0-15%), the leading, fast leg generated a large peak negative peak in power while the trailing, slow leg generated a relatively smaller positive peak. This relationship reversed during the later portion of the stride cycle (∼40-70% of the gait cycle) such that the fast leg generated a burst of positive power while the slow leg generated a smaller burst of negative power. Each trace is an average of all participants (n = 16). The stride cycle begins and ends at foot-strike of the fast limb. Blue, yellow, and red traces correspond to split-belt walking at step length asymmetries of -15%, 0%, and +15%, respectively.

**Figure 5:**
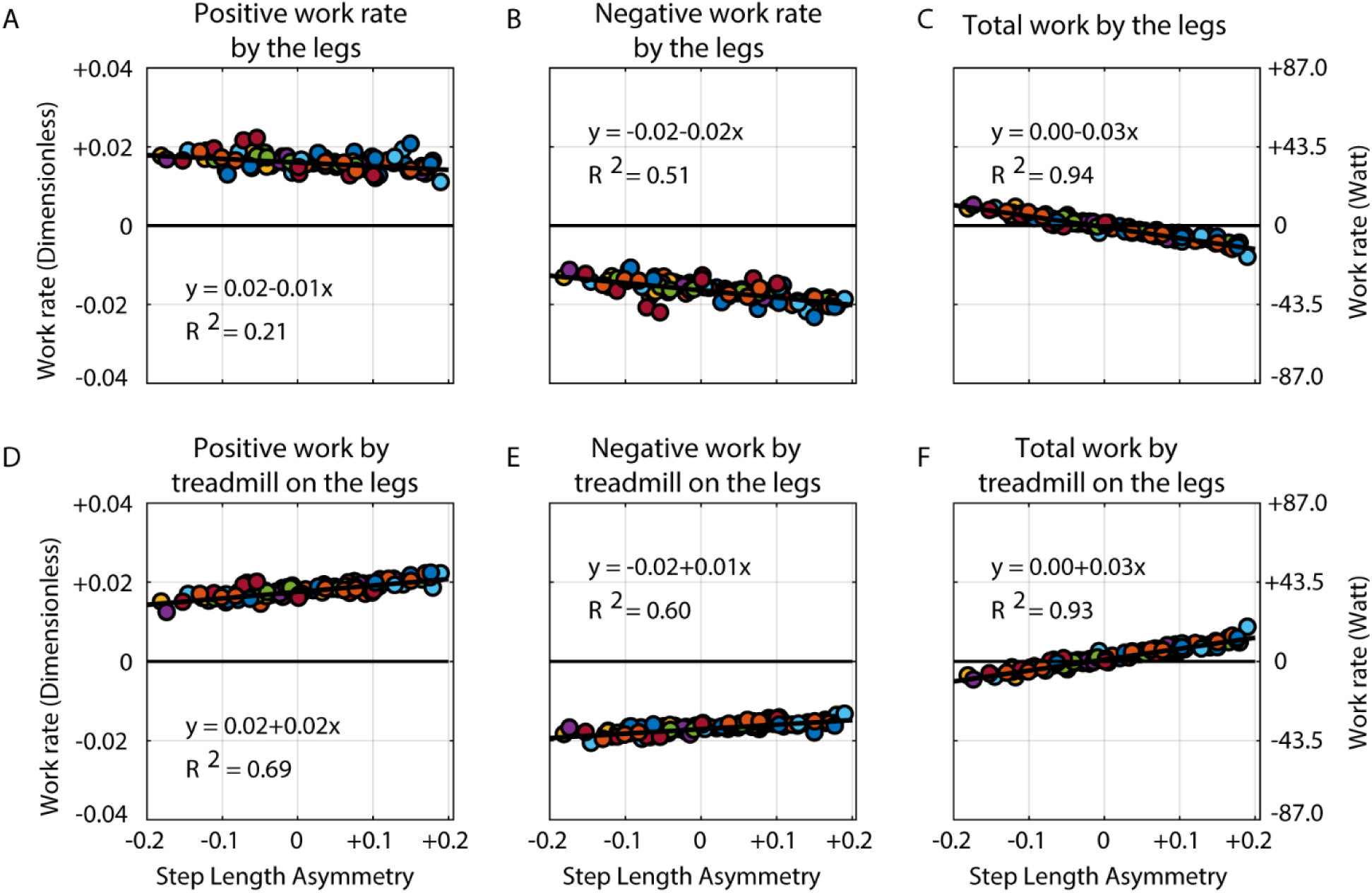
Average rate of work performed by the legs and the treadmill belts as a function of step length asymmetry. A) Positive work performed by the legs, B) negative work performed by the legs, and C) total work performed by the legs. D) Positive work performed on the body by the treadmill, E) negative work performed on the body by the treadmill, and F) total work performed by the treadmill on the body. Work rate is expressed in dimensionless units on the left y-axis and in Watts on the right y-axis. Each data point represents a single trial for an individual participant and each color represents a different participant.

We predicted that the treadmill would act as an assistive device at positive step length asymmetries by performing net positive work on the body, and our observations were consistent with this prediction (Figure 4C). As asymmetry became more positive, there was a ∼28% increase in the amount of positive work performed by the belts on the body (Figure 5D, β =0.016, 95% CI [0.014, 0.018], p=1.64e-29) and a ∼15% decrease in the amount of negative work performed by the belts on the body (Figure 5E, β =0.011, 95% CI [0.010, 0.013], p=1.58e-23). Together, these changes resulted in a shift from the treadmill performing mostly negative work on the body and extracting energy from the person at negative asymmetries to performing mostly positive work on the body and adding energy to the person at positive asymmetries (Figure 5F, β =0.028, 95% CI [0.026, 0.029], p=4.76e-67).

The legs’ shift toward performing net negative work at positive step length asymmetries was primarily driven by changes in the fast leg. Across participants, we observed both a ∼30 % reduction in positive work (Figure 6A, β =0.010, 95% CI [-0.012, 0.008], p=7.34e-17) and a ∼36% increase in negative work performed by the fast leg (Figure 6C, β = -0.012, 95% CI [-0.014, -0.009], p=7.63e-14) at increasingly positive asymmetries. In contrast, there were no changes in positive work performed by the leg on the slow belt (Figure 6B, p=0.53), but there was a ∼21% increase in negative work by the slow leg at positive asymmetries (Figure 6D, β = -0.007, 95% CI [-0.010, -0.004], p=4e-5).

**Figure 6:**
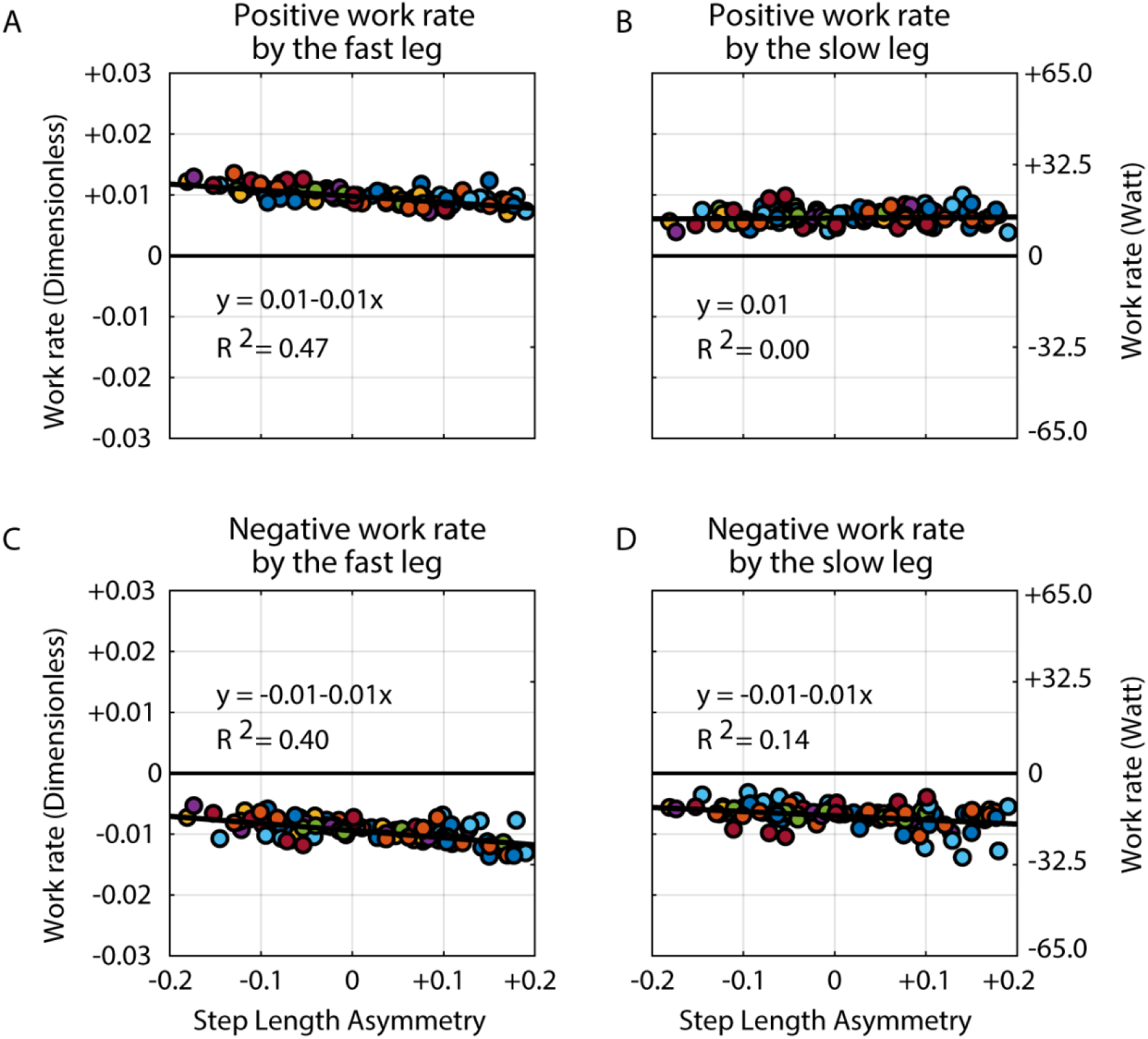
Average rate of work performed by the individual legs across levels of step length asymmetry. A) Positive work performed by the fast leg, B) positive work performed by the slow leg, C) negative work performed by the fast leg, and D) negative work performed by the slow leg. Work rate is expressed in dimensionless units on the left y-axis and in Watts on the right y-axis. Each data point represents a single trial for an individual participant and each color represents a different participant.

As the treadmill performed increasingly more positive work on the body, there was a proportional reduction in positive work performed by the legs (Figure 7, β = -0.33, 95% CI [-0.43, -0.23], p=2.8e-9). The slope of this relationship suggests that participants exploited the work performed by the treadmill belts with an effectiveness of approximately 33%. That is, for every 3 Joules of positive work performed by the belts on the person, the person reduced the positive work performed by the legs by about 1 Joule.

**Figure 7:**
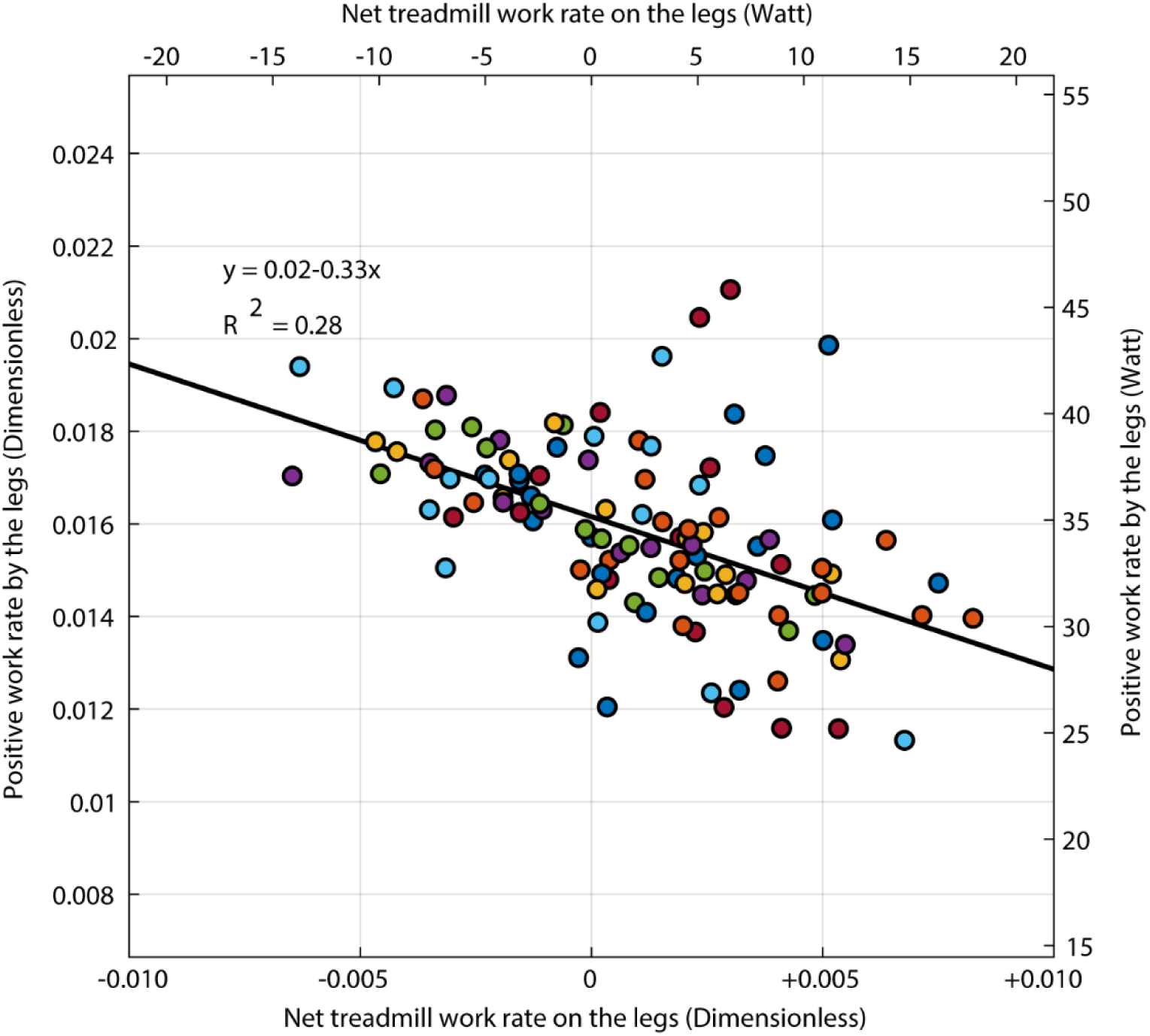
Relationship between the average rate of positive work performed by the legs and the average rate of work performed by the treadmill on the body. Work rate is expressed in dimensionless units on the lower x-axis and the left y-axis and in Watts on the upper x-axis and right y-axis. Each data point represents a single trial for an individual participant and each color represents a different participant.

### Assistance Provided by the Treadmill Led to a Reduction in Metabolic Cost

The assistance provided by the treadmill at positive step length asymmetries was not only associated with a reduction in positive work performed by the legs, but was also associated with a reduction in metabolic cost (Figure 8A, β = -1.54, 95% CI [-2.07, -1.02], p=5.34e-8). Since the assistance provided by the treadmill was associated with a reduction in positive work performed by the legs, we also found that metabolic power was strongly correlated with the total positive work performed by the legs (Figure 8B, β = 0.105, 95% CI [0.091, - 0.119], p=1.89e-27). The slope of this relationship suggests that positive work was performed at an efficiency of about 10%. Overall, by exploiting the assistance provided by the treadmills, participants achieved an approximately 14% reduction (8 to 18% reduction, 95% CI) in metabolic cost relative to the most costly level of step length asymmetry (Figure 8C).

**Figure 8:**
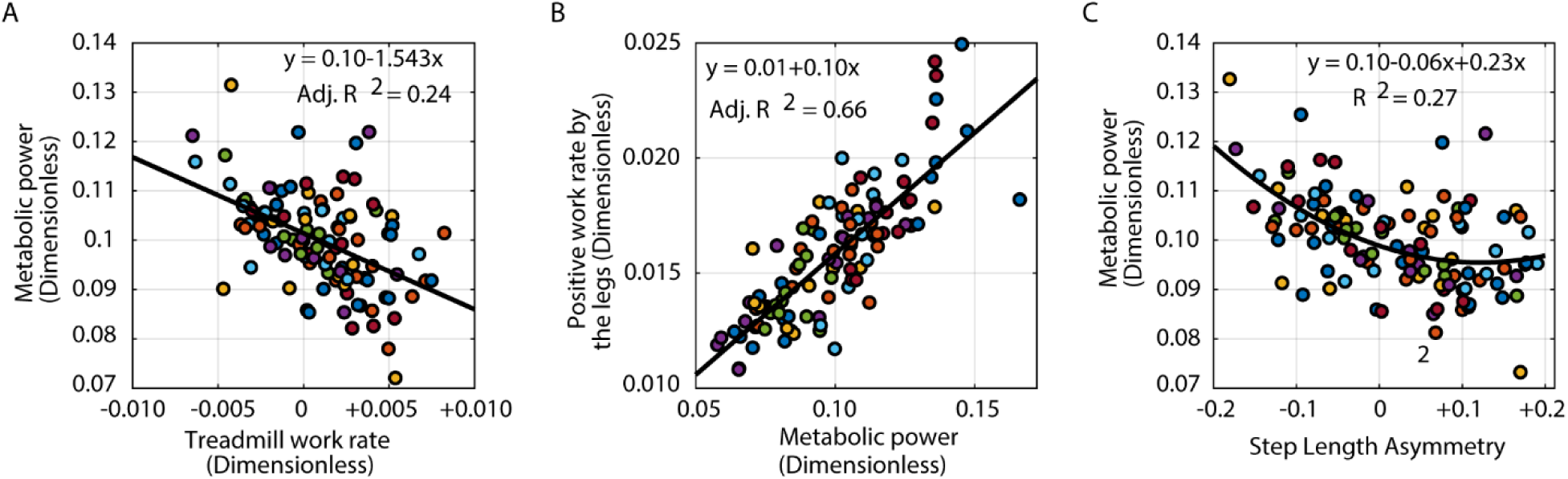
Metabolic power, mechanical work, and asymmetry. A) Metabolic power as a function of the rate of work performed by the treadmill on the body. B) Positive work rate versus metabolic power. C) Metabolic power as a function of step length asymmetry. Each data point represents a single trial for an individual participant.

### The Ability to Use Step Length Asymmetry to Exploit Assistance Provided by the Treadmill is Bounded

Although the increase in positive work performed by the treadmill led to a reduction in metabolic cost, we also wanted to determine if this ability to exploit the work performed by the treadmill increased indefinitely or whether there was a specific level of asymmetry that minimized metabolic cost. Our results supported the existence of an energetically optimal level of step length asymmetry as a regression model including both linear and quadratic terms explained the relationship between metabolic cost and step length asymmetry better than a simple linear model (LRStat = 4.83, p=0.028, adjusted R^2^ = 0.27, Figure 8C). Bootstrap analyses indicated that the asymmetry that minimized metabolic cost had a 95% confidence interval of 0.06 and 0.38, which is consistent with our prediction that positive asymmetries minimize energetic cost.

Although metabolic cost generally decreased with increasingly positive asymmetries, we expected that it would increase again at large positive asymmetries. We confirmed this finding in two participants, who each completed two additional trials where they walked at target levels of 0.20 and 0.25 step length asymmetry. The metabolic cost for three out of four of these trials was 0.027 ± 0.022 greater than the minimum cost predicted from the regression fit. These increases in metabolic cost at extreme positive asymmetries support the quadratic relationship that we determined in our analysis.

### Participants Chose Positive Step Length Asymmetries When Allowed to Freely Adapt

Lastly, we found that participants chose to walk at a level of step length asymmetry near that which minimized metabolic cost when they were allowed to freely select their walking pattern. After experiencing a range of asymmetries during the cost mapping trials, participants tended to plateau at step length asymmetries that were more positive than their natural baseline asymmetry (Figure 9, mean: 0.028, 95% CI [0.0018, 0.0434], p=0.010) though they had shorter stride lengths during adaptation (95% CI [929 mm, 999 mm]) than they did when walking at similar levels of asymmetry during the split-belt trials with visual feedback (95% CI [47, 93] mm shorter strides during adaptation, p= 1.03e-05). This step length asymmetry markedly differs from the behavior typically observed within a single session of adaptation where participants most often plateau at slightly negative asymmetries [28,29]. In addition, there was no significant difference between the metabolic cost measured at the end of the adaptation trial (0.086, 95% CI [0.074, 0.10]) and the minimum predicted from the regression fit (0.096, p=0.12).

**Figure 9:**
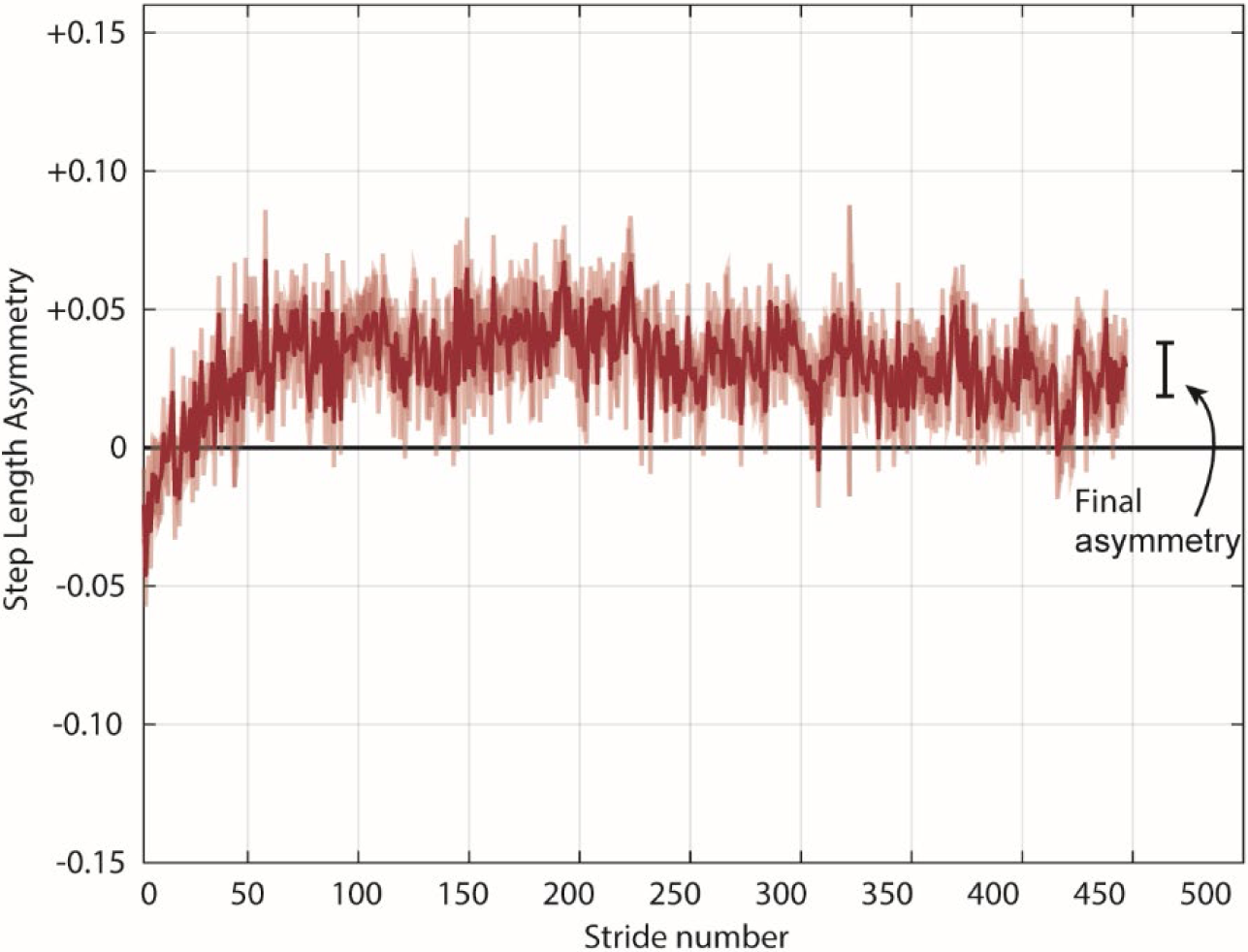
Adaptation of step length asymmetry during the adaptation trial in the absence of visual feedback. Participants consistently selected positive asymmetries near those which minimized metabolic cost during the cost mapping trials. The data are an average of all participants and the shaded area around the average trace represents the standard error. The final error represents the average ± standard error level of asymmetry at the end of the split-belt adaptation trial.

## Discussion

Learning to gain assistance from external sources is a general problem for the nervous system. We explored this problem using a split-belt treadmill paradigm to determine whether people learn to harness energy from the differences in belt speeds to reduce the metabolic cost of walking. People exploited the assistance provided by the treadmill by changing their step lengths such that the treadmill performed net positive work on the body, thereby allowing the legs to perform net negative work. This shift toward performance of negative work by the legs was associated with a reduction in metabolic cost which likely reflects the energetic benefits of negative work [30]. Therefore, the reductions in asymmetry commonly observed during split-belt walking can be interpreted as a strategy generated by the neuromotor system to take advantage of the work performed by the treadmill to reduce energetic cost [19,21].

Similar to using a powered exoskeleton, individuals can learn to coordinate their movement to maximize the assistance from the treadmill’s motors. We find that individuals use the assistance from the split-belt treadmill with an effectiveness of 33%, i.e. they reduce their positive work by 1 J for every 3 J of positive work done by the treadmill. While this is not a commonly reported metric in exoskeleton use, Sawicki and Ferris [10] used a similar metric to show that their ankle exoskeletons reduced joint mechanical power by 41% relative to the mechanical power provided by their ankle exoskeleton—similar to our results. We focus on this metric because while the maximum effectiveness possible in powered exoskeletons may be system-dependant and hard to quantify, we can provide a reasoned estimate of the maximum possible effectiveness for split-belt treadmill walking. When the treadmill does positive work on the person, it applies a negative force on the person that has to be cancelled out by an equal positive propulsive force applied by the trailing leg on the treadmill. In applying this propulsive force, the trailing leg has to perform positive work on the treadmill. In fact, the minimum amount of positive work necessary depends on the ratio of the belt speeds. In our experiment, the fast belt moved three times faster than the slow belt. Since work is the time integral of the dot product of force and velocity, this means that the trailing leg on the slow belt has to at least perform positive work that is roughly one third of the negative work performed by the leading leg on the fast belt. This means that the participants in our study could have at most achieved an effectiveness of ∼67%. We only include the fast step in this calculation because any braking force on the slow belt, only further decreases this value.

It is possible that individuals cannot, in practice, exploit the positive work from the treadmill to the degree we have suggested here. One strategy to improve effectiveness in split-belt walking is for individuals to use more positive step length asymmetries. However, the extreme positive asymmetries needed to maximize the positive work performed by the treadmill likely challenge anatomical constraints. In addition, even at less extreme positive asymmetries, an individual might choose to perform more positive work, and consequently also negative work, than necessary simply to feel safe in the new gait [31]. We suspect that with practice and appropriate guidance, the effectiveness observed in our study can be improved from that observed here. This is one of the goals of our future research.

The reduction in positive mechanical work we observed was accompanied by a reduction in metabolic cost of 14%. This reduction is comparable to that observed in powered lower-limb exoskeletons that can currently achieve reductions up to 17% [32]. Both effectiveness and energetic benefits incurred as a result of learning to walk in lower-limb exoskeletons are similar to that observed in split-belt walking. As described earlier, we suspect that people can be taught to improve their effectiveness and maximize energetic benefits in split-belt walking, and also when using powered exoskeletons. Given the convenience of acquiring and operating a split-belt treadmill, it makes a good paradigm to understand how the human nervous system learns to walk in novel systems, and thus improve the benefits obtained by individuals while walking with exoskeletons.

Traditionally, studies of split-belt adaptation have shown that after 10-20 minutes of adaptation, individuals converge to step length asymmetries near zero [15,19]. This adaptation toward a step length asymmetry of zero is consistent with the hypothesis that step length asymmetry is treated as an error by the nervous system. In this hypothesis, the difference between expected and achieved sensory feedback during movement, known as sensory prediction error, drives motor adaptation [33–35]. Reductions in step length asymmetry are also consistent with the hypothesis that individuals converge towards habitual behaviors when exposed to novel environments [36,37]. Based on this hypothesis, people adopt steps of equal length because this is the habitual pattern they use regularly. One challenge to both of these notions is that people adopt asymmetric step times in order to take steps of equal length [38] and thus, it is not immediately apparent why the nervous system would choose to reduce errors in step length but not in time. In addition, these asymmetries in step time could lead to sensory prediction errors and are non-habitual behaviors in the time domain. Further evidence that adaptation is not purely driven by sensory prediction errors related to step length asymmetry was provided by a recent study showing that sensory recalibration and motor recalibration have different timescales [39]. In this study, the authors postulated that if we recalibrate motor commands in response to sensory prediction errors, then error perception is also updated [40]. Therefore, if the same neural processes drive motor and perceptual recalibration during locomotor adaptation, they would change over a similar timescale. However, the authors found that motor and perceptual adaptation to differences in belt speeds occurred over different timescales and are likely independent of each other. The authors also found that after multiple days of adaptation, individuals plateau at positive step length asymmetries. These results, together with our findings that individuals adopt positive asymmetries after being exposed to the cost landscape, refute the idea that split-belt adaptation is explained by the nervous system’s desire to minimize perceived errors in step length asymmetry or converge towards habitual behaviors. Instead, energy optimization explains both why people reduce step length asymmetry during single sessions of split-belt adaptation and why they adopt positive asymmetries when provided with more extensive experience.

A logical follow-up question is, if energetic optimization is the goal of split-belt adaptation, why are positive asymmetries that minimize mechanical work and metabolic cost not observed during adaptation? One potential explanation is that the energetic savings for positive asymmetries are minor compared to the cost of symmetry, and this might impede the optimization process. Given that the confidence interval for the step length asymmetry associated with the lowest metabolic cost ranged from 6 to 38%, the energetic gradient might be too shallow for people to obtain meaningful energetic reductions from walking with positive asymmetries. In fact, our results show that the optimal step length asymmetry reduced metabolic cost by only 2% compared to symmetry. Despite these small savings, after exposure to the step length asymmetry landscape, participants in our study plateaued at positive asymmetries during adaptation. This suggests that people may be willing to adjust how they walk for savings of less than 5% as reported in previous work [20].

Alternatively, positive asymmetries may not have been observed during typical locomotor adaptation studies because energetic optimization occurs over a timescale that is longer than that commonly used in adaptation studies [9, 10]. To date, most locomotor adaptation studies have been performed using single session paradigms of 10 to 20 minutes in duration [15,19]. Thus, short, single bout studies may not provide enough time or experience for individuals to fine tune their steps lengths to achieve the more energetically optimal positive asymmetries. Consistent with the interpretation that energy optimization occurs over a longer timescale, people tend to reach positive asymmetries when allowed to adapt to a split-belt treadmill over multiple days [39]. Surprisingly, the visual feedback in our experiment, which exposed participants to positive asymmetries, accelerated the convergence towards positive asymmetries in a single session to that which occurs during multi-day adaptation. The longer timescale for energetic optimization is further supported by work in the upper extremity, which shows that improvements in task performance and fine tuning of upper extremity muscle activation, occurred over a faster timescale than energy minimization [25,41]. Overall, we conclude that the symmetric steps commonly observed at the conclusion of previous split-belt adaptation studies and the associated reductions in energetic cost [18,19], may be only a partial picture of a slower energetic optimization process that plateaus at positive asymmetries.

One of the features of our study is that we constrained stride lengths to those measured during baseline with the belts tied at 1m/s. Given this constraint, the metabolic optima that we found in this study is a local minima for that specific stride length, and may not be a global minimum. We imposed this constraint as it would not be practical to characterize the metabolic cost landscape across both the dimensions of step length asymmetry and stride length. Moreover, post-hoc analysis of data from a previous adaptation study [21] showed that there was no change in stride length during split-belt adaptation compared to baseline walking (p=0.389). Our constraint on stride length is particularly relevant given the results of the adaptation trial in the current study, where 15/16 participants adapted to the split-belt treadmill using shorter stride lengths. Whether optimization of stride length can further reduce positive work and metabolic cost during locomotor adaptation remains to be seen.

In conclusion, the process by which people adapt to walking on a split-belt treadmill is just one example of a broad class of tasks in which the nervous system learns to exploit external assistance to improve economy. A common feature of these types of tasks is that the process of optimizing the use of assistance may proceed quite gradually in the absence of guided experience. Ultimately, understanding how best to guide people through a range of experiences capable of accelerating the learning and optimization process has important implications for maximizing the utility of assistive devices such as exoskeletons and prostheses.

## Materials and Methods

### Experiment Protocol

Sixteen healthy participants (7 female, 9 male, age 27 +/-3.5 years) completed our study. All experimental procedures were approved by the University of Southern California Institutional Review Board and each participant provided written informed consent before testing began. All aspects of the study conformed to the principles described in the Declaration of Helsinki.

We used a biofeedback-based protocol to map the relationship between mechanical work, metabolic cost, and step length asymmetry during split-belt walking (Figure 2B). Participants first walked on an instrumented split-belt treadmill (Fully Instrumented Treadmill, Bertec Corporation, OH) with both belts at 1.0 m/s while we measured their baseline step lengths and metabolic cost. In the next trial, we introduced the visual feedback and participants walked with both belts at 1.0 m/s, while matching their step lengths to the visual targets (see below). For the next seven trials, we set the belt speeds to a 3:1 ratio, with the left belt at 1.5 m/s and the right belt at 0.5 m/s. Participants performed the split-belt trials with visual feedback providing target step length asymmetries of 0.00, +/-0.05, +/-0.10 and +/-0.15 (see below). These asymmetries were relative to each participant’s baseline and we presented these trials in random order. To determine whether mechanical work or energetic cost increased at extreme positive asymmetries, two participants performed two additional trials at asymmetries of +0.20 and +0.25. Each of these trials was six minutes in duration.

After the visual feedback trials, participants completed a 10-minute split-belt adaptation trial with no visual feedback of their step lengths and no explicit instructions of the step lengths they should maintain. The purpose of the final trial was to test whether participants would converge towards the metabolically or mechanically optimal level of asymmetry observed during the previous split-belt trials. During all walking trials, participants wore a harness designed to prevent falls while providing no body weight support. Participants did not hold on to handrails during any portion of the trials. After each walking trial, participants sat down and rested for at least four minutes, and we visually inspected measured metabolic cost to ensure that it had returned to resting levels before beginning the next trial.

For all feedback conditions, participants relied on a monitor located at eye-level in front of the treadmill to target the desired step length targets. We measured the position of ankle markers placed bilaterally on the lateral malleoli at 100 Hz using an 11 camera Qualisys Oqus camera system (QTM, Sweden) and the visual display was controlled by custom software written in Vizard (Worldviz, Santa Barbara, CA). A fourth-order low-pass digital Butterworth filter smoothed marker data using a cut-off frequency of 10 Hz. We defined peak fore-aft position of the filtered ankle marker trajectories as heel-strike, and step length as the distance between right and left ankle markers at this instance. The monitor displayed the ankle position for the intervals where the ground reaction forces were less than 20N [42], which corresponded to swing phase. At foot-strike, defined as the point when the ground reaction force exceeded 20N, the ankle marker disappeared from the visual feedback. We instructed participants to step such that the location of the ankle marker at heel-strike would match the top a vertical bar in the visual display (Figure 2B). Participants targeted the left bar for the left leg and the right bar for the right leg. We constrained the sum of the step lengths to be equal to the baseline stride length (Equation 1), therefore participants adjusted the individual step lengths while maintaining stride length equal to that measured during the baseline trial. Note that the baseline speed was 1m/s, which is the average of the split-belts speed:

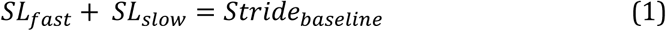

We constrainined the stride length so that the desired step lengths would be fixed for a given level of step length asymmetry. To reinforce performance, the display provided participants with a score for each step at heel-strike, according to the following equation, rounded to the nearest integer:

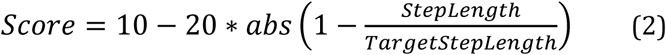

Participants only received a score if their step lengths were within eight standard deviations from the target. For example, for a representative participant, the average step length measured on the baseline trial was 560mm, thus, the achieved step length must be within 14 mm of the target to obtain a score of 10 on each side. For this same participant, the standard deviation was 17mm. If the step length was off-target by more than 136 mm, the participant did not receive a score. We verbally encouraged participants to obtain the maximum score of 10 points for all steps.

Each participant’s baseline asymmetry was defined as a step length asymmetry of zero, and all levels of asymmetry were expressed relative to this baseline. As per convention [19,21], we defined step length asymmetry as:

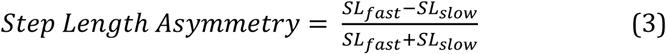

Here *SL*_*fast*_ is the step length at the instant the leading leg heel-strikes on the fast belt, and *SL*_*slow*_ is the step length at the instant the leading leg heel-strikes on the slow belt. Negative values correspond to longer steps with the slow (right) leg and positive values correspond to longer steps with the fast (left) leg. Accordingly, we use the last 100 strides of each trial to obtain the average step length asymmetry.

There may be more direct methods to change leg coordination to gain positive work from the treadmill than manipulating step length asymmetry. We choose to use step length asymmetry for two reasons. First, prior research has shown that during split-belt walking, people increase fast step lengths by stepping further forward on the fast belt and decrease slow step length by lifting the trailing fast leg sooner, both during adaptation and when increasingly more positive asymmetries are enforced with visual feedback [15,21,43,44]. Second, the literature on adapting to split-belts is primarily focused on considering adaptation as a process that minimizes step length asymmetry error [33,44] and keeping our manipulation in terms of step length asymmetry helps make clear that there are alternative explanations for why the nervous system may adapt to reduce negative step length asymmetries. We anticipate that stepping further forward on the fast belt will lead to more braking force by the fast leg around heel-strike, and thus more positive work on the person by the belt. We also anticipate that short steps with the slow leg will be associated with reduced propulsive force generated by the fast leg around toe-off, and thus less positive work generated by that leg. In addition, longer steps with the fast belt will also lead the slow leg to lift off at a greater hip angle, generating increased propulsion.

### Analysis

We assessed metabolic cost by determining the rates of oxygen consumption (*V*_*O2*_) and carbon dioxide production (V_CO2_) using a TrueOne^®^ 2400 system (Parvomedics, UT). The metabolic cart recorded data on a breath-by-breath basis and subsequently, we re-sampled these data at a frequency of 0.1 Hz and averaged for smoothing in 10s bins. Since it takes approximately three minutes for oxygen consumption and carbon dioxide production by the body to reach steady-state in a task, we first identified the time point closest to the third minute of the trial [20]. We then measured the total V_O2_ consumed and the V_CO2_ produced from that time point onwards until the end of the trial. We then estimated the energy consumed during the last three minutes using the standard Brockway equation [27] as follows:

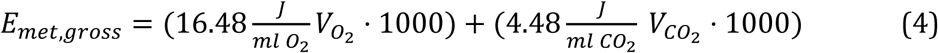

From here, we dived *E*_*met,gross*_ by the exact duration (T) over which it was calculated to obtain an estimate of the gross metabolic rate *P*_*met,gross*_ measured in Watts. Finally, we subtracted each participants’ standing metabolic rate from each walking trials. Thus, all metabolic rate values presented here are net metabolic rate.

We estimated the mechanical work performed by the legs using an extension of the individual limbs method [18,26]. This method approximates the legs as massless pistons and the entire body as a point mass acting at the center of mass (Figure 1A). We use a reference frame attached to the stationary ground—the belt speeds and center of mass velocity are relative to this reference frame. We measured individual leg ground reaction forces from the instrumented treadmill at 1,000 Hz and filtered this signal with 20 Hz cut-off low-pass zero-lag digital Butterworth filter. We segmented the ground reaction forces into strides using a vertical ground reaction force threshold of 32N [16] to identify the beginning and end of each stride, and performed the following analysis on a stride-by-stride basis.

We calculated the medio-lateral, fore-aft and vertical center of mass velocities by first calculating the center of mass accelerations as the sum of the forces acting on the body normalized for body mass [18,26]. We estimated body mass as the average vertical force during the final 100 strides of the trial, divided by the acceleration due to gravity (9.81 m/s^2^). We calculated center of mass velocities from the time integral of the center of mass accelerations. We determined the integration constants by requiring the average center of mass velocity over a stride to be zero in each direction because net movement in any direction must on average be small on a treadmill.

The total mechanical power generated by each leg is composed of power generated by the leg on the body and the power generated by the leg on the treadmill’s belts. We define the mechanical power generated on the body by a leg as the dot product of the ground reaction force from that leg and the center of mass velocity. Similarly, we define the mechanical power generated by a leg on a belt as the dot product of the force generated by the leg on the belt, which is equal and opposite to the ground reaction force measured by the treadmill, and the velocity of the corresponding belt. The medio-lateral and vertical components of belt velocity are zero, and the fore-aft component is either -0.5 m/s (slow belt), -1.5 m/s (fast belt), or -1.0 m/s (tied belts). For each leg, we then calculated the instantaneous sum of the two powers to obtain the total instantaneous mechanical power generated by that leg (Figure 1B). We calculate the instantaneous power generated by the slow and fast belts on the body as the dot product of the ground reaction force measured from that belt with that belt’s velocity.

To determine the total positive and negative work performed by a leg or a belt, we calculated the time integral of the positive or negative portion of its instantaneous power over the stride cycle [45]. The net work performed by a leg or a belt is the time integral of the full instantaneous power over the stride cycle. We express all measures of work as work rates by dividing each measure by stride duration. As with step length asymmetry, measures of work rate for each trial are the average values over the last 100 strides.

We converted the metabolic rate and mechanical work rate to dimensionless units to reduce variability between subjects. We divided each individual’s values by *ml*^*0.5*^*g*^*1.5*^ here m is their body mass, *l* is their leg length and *g* is gravity (9.81 m/s^2). Thus, using the average body mass (75.4 kg) and average leg length (0.88 m) of our participants, all reported dimensionless values can be approximately redimensionalized to the average values in Watts by multiplying by 2.17 x 10^3^.

### Statistical Analyses

A participant’s mechanical work and metabolic cost depend not only on the conditions that we control but also on differences between individuals. Our purpose here is to test predictions about the former—we do not, for example, seek to explain the differences in metabolic cost between individuals for a given condition. Consequently, we used mixed-effect regression models that allowed individualized intercepts but shared a fixed dependence on the independent variables of interest. These models captured the relationship between 1) measure of foot placement and step length asymmetry, 2) measures of mechanical work and step length asymmetry, 3) measures of mechanical work performed by the legs and the mechanical work performed by the treadmill, 4) metabolic power and mechanical work performed by the treadmill, and 5) metabolic power and step length asymmetry. All models included a random intercept for each participant to account for unknown, subject-specific effects. We used a modified version of the marginal R^2^ for linear mixed effect models [46] to compute the variance explained by the fixed components of our linear models. We computed R^2^ as the ratio of the variance computed from the fixed effects and the sum of the variance from both the fixed effects and residuals from the regression model. We used this approach in lieu of the conditional R^2^, which accounts for the variance explained by both the fixed and random effects, because we were only interested in quantifying the explanatory value of the fixed effects. To make our figures consistent with our statistical analysis approach, we removed the individualized intercepts from each participant’s data before generating each scatterplot. Lastly, we determined if participants plateaued at a step length asymmetry that differed from baseline during the adaptation period using a paired-samples t-test. We conducted all statistical analyses in Matlab R2017a (Mathworks, Natick, MA) and set statistical significance level to p < 0.05.

## Authorship Contributions

Conceptualization, J.M.F., J.M.D, S.N.S, and N.S; Methodology, J.M.F, J.M.D, S.N.S, and N.S; Investigation, N.S.; Formal Analysis and Data Curation – J.M.F., J.M.D, S.N.S, and N.S; Writing – Original Draft, J.M.F, J.M.D, S.N.S, and N.S.; Writing–Review & Editing, J.M.F, J.M.D, S.N.S, and N.S.; Visualization:.M.F, J.M.D, S.N.S, and N.S; Funding Acquisition, J.M.F and N.S.; Resources, J.M.F.; Supervision, J.M.F and J.M.D.

## References

1. McGeer T. Passive Dynamic Walking. Int J Rob Res. 1990;9: 62–82.

2. Mochon S, McMahon TA. Ballistic walking: an improved model. Math Biosci. 1980;52: 241–260. doi:10.1016/0025-5564(80)90070-X

3. Herr H. Exoskeletons and orthoses: Classification, design challenges and future directions. J Neuroeng Rehabil. 2009;6: 1–9. doi:10.1186/1743-0003-6-21

4. Ferris DP. The exoskeletons are here. J Neuroeng Rehabil. 2009;6: 1–3. doi:10.1186/1743-0003-6-17

5. Ferris DP, Sawicki GS, Daley MA. A Physiologist’s Perspective on Robotic Exoskeletons for Human Locomotion. Int J Humanoid Robot. 2007;04: 507–528. doi:10.1142/S0219843607001138

6. Gordon KE, Ferris DP. Learning to walk with a robotic ankle exoskeleton. J Biomech. 2007;40: 2636–2644. doi:10.1016/j.jbiomech.2006.12.006

7. Gordon KE, Kinnaird CR, Ferris DP. Locomotor adaptation to a soleus EMG-controlled antagonistic exoskeleton. J Neurophysiol. 2013;109: 1804–14. doi:10.1152/jn.01128.2011

8. Koller JR, Jacobs DA, Ferris DP, Remy CD. Learning to walk with an adaptive gain proportional myoelectric controller for a robotic ankle exoskeleton. J Neuroeng Rehabil. Journal of NeuroEngineering and Rehabilitation; 2015;12: 1–14. doi:10.1186/s12984-015-0086-5

9. Selinger JC, Donelan JM. Myoelectric Control for Adaptable Biomechanical Energy Harvesting. IEEE Trans Neural Syst Rehabil Eng. IEEE; 2016;24: 364–373. doi:10.1109/TNSRE.2015.2510546

10. Sawicki GS, Ferris DP. Mechanics and energetics of level walking with powered ankle exoskeletons. J Exp Biol. 2008;211: 1402–1413. doi:10.1242/jeb.009241

11. Malcolm P, Derave W, Galle S, De Clercq D. A Simple Exoskeleton That Assists Plantarflexion Can Reduce the Metabolic Cost of Human Walking. PLoS One. 2013;8: 1–7. doi:10.1371/journal.pone.0056137

12. Galle S, Malcolm P, Collins SH, De Clercq D. Reducing the metabolic cost of walking with an ankle exoskeleton: interaction between actuation timing and power. J Neuroeng Rehabil. Journal of NeuroEngineering and Rehabilitation; 2017;14: 1–16. doi:10.1186/s12984-017-0235-0

13. Jackson RW, Collins SH. An experimental comparison of the relative benefits of work and torque assistance in ankle exoskeletons. J Appl Physiol. 2015;119: 541–557. doi:10.1152/japplphysiol.01133.2014

14. Zhang J, Fiers P, Witte KA, Jackson RW, Poggensee KL, Atkeson CG, et al. Human-in-the-loop optimization of exoskeleton assistance during walking. ScienceScience. 2017;356: 1280–1284. doi:10.1126/science.aal5054

15. Reisman DS, Block HJ, Bastian AJ. Interlimb coordination during locomotion: what can be adapted and stored? J Neurophysiol. 2005;94: 2403–2415. doi:10.1152/jn.00089.2005

16. Dietz V, Zijlstra W, Duysens J. Human neuronal interlimb coordination during split-belt locomotion. Exp Brain Res. 1994;101: 513–520. doi:10.1007/BF00227344

17. Prokop T, Berger W, Zijlstra W, Dietz V. Adaptational and learning processes during human split-belt locomotion: interaction between central mechanisms and afferent input. Exp Brain Res. 1995;106: 449–456.

18. Selgrade BP, Thajchayapong M, Lee GE, Toney ME, Chang Y-H. Changes in mechanical work during neural adaptation to asymmetric locomotion. J Exp Biol. 2017;9993: jeb.149450. doi:10.1242/jeb.149450

19. Finley JM, Bastian AJ, Gottschall JS. Learning to be economical: the energy cost of walking tracks motor adaptation. J Physiol. 2013;591: 1081–1095. doi:10.1113/jphysiol.2012.245506

20. Selinger JC, O’Connor SM, Wong JD, Donelan JM. Humans Can Continuously Optimize Energetic Cost during Walking. Curr Biol. Elsevier Ltd; 2015;25: 1–5. doi:10.1016/j.cub.2015.08.016

21. Sánchez N, Park S, Finley JM. Evidence of Energetic Optimization during Adaptation Differs for Metabolic, Mechanical, and Perceptual Estimates of Energetic Cost. Sci Rep. 2017;7. doi:10.1038/s41598-017-08147-y

22. Burdett RG, Skrinar GS, Simon SR. Comparison of mechanical work and metabolic energy consumption during normal gait. J Orthop Res. 1983;1: 63–72. doi:10.1002/jor.1100010109

23. Umberger BR, Gerritsen KGM, Martin PE. A Model of Human Muscle Energy Expenditure. 2003;6: 99–111. doi:10.1080/1025584031000091678

24. Doke J, Donelan JM, Kuo AD. Mechanics and energetics of swinging the human leg. J Exp Biol. 2005;208: 439–445. doi:10.1242/jeb.01408

25. Huang HJ, Kram R, Ahmed AA. Reduction of metabolic cost during motor learning of arm reaching dynamics. J Neurosci. 2012;32: 2182–2190. doi:10.1523/JNEUROSCI.4003-11.2012

26. Donelan JMM, Kram R, Kuo AD. Simultaneous positive and negative external mechanical work in human walking. J Biomech. 2002;35: 117–124. doi:10.1016/S0021-9290(01)00169-5

27. Brockway JM. Derivation of formulae used to calculate energy expenditure in man. Hum Nutr Clin Nutr. 1987;41: 463–71.

28. Leech KA, Day KA, Roemmich RT, Bastian AJ. Movement and perception recalibrate differently across multiple days of locomotor learning. J Physiol. 2018;

29. Leech KA, Roemmich RT, Bastian AJ. Creating flexible motor memories in human walking. Sci Rep. Springer US; 2018;8: 1–10. doi:10.1038/s41598-017-18538-w

30. Herzog W. The mysteries of eccentric muscle action. J Sport Heal Sci. Elsevier B.V.; 2018;00: 5–6. doi:10.1016/j.jshs.2018.05.006

31. Hunter LC, Hendrix EC, Dean JC. The cost of walking downhill: Is the preferred gait energetically optimal? J Biomech. 2010;43: 1910–1915. doi:10.1016/j.jbiomech.2010.03.030

32. Ding Y, Kim M, Kuindersma S, Walsh CJ. Human-in-the-loop optimization of hip assistance with a soft exosuit during walking. Sci Robot. 2018;3: eaar5438. doi:10.1111/irfi.12020

33. Roemmich RT, Long AW, Bastian AJ. Seeing the Errors You Feel Enhances Locomotor Performance but Not Learning. Curr Biol. Elsevier Ltd.; 2016;26: 2707–2716. doi:10.1016/j.cub.2016.08.012

34. Vazquez A, Statton MA, Busgang SA, Bastian AJ. Split-belt walking adaptation recalibrates sensorimotor estimates of leg speed but not position or force. J Neurophysiol. 2015;114: 3255–3267. doi:10.1152/jn.00302.2015

35. Shadmehr R, Smith M a, Krakauer JW. Error correction, sensory prediction, and adaptation in motor control. Annu Rev Neurosci. 2010;33: 89–108. doi:10.1146/annurev-neuro-060909-153135

36. de Rugy a., Loeb GE, Carroll TJ. Muscle Coordination Is Habitual Rather than Optimal. J Neurosci. 2012;32: 7384–7391. doi:10.1523/JNEUROSCI.5792-11.2012

37. Loeb GE. Optimal isn’t good enough. Biol Cybern. 2012;106: 757–765. doi:10.1007/s00422-012-0514-6

38. Finley JM, Long A, Bastian AJ, Torres-Oviedo G. Spatial and Temporal Control Contribute to Step Length Asymmetry During Split-Belt Adaptation and Hemiparetic Gait. Neurorehabil Neural Repair. 2015; doi:10.1177/1545968314567149

39. Leech KA, Roemmich RT, Day KA, Bastian AJ. Movement and perception recalibrate differently across multiple days of locomotor learning. J Neurophysiol. 2018;114: 608–623. doi:10.1152/jn.00628.2014

40. Izawa J, Rane T, Donchin O, Shadmehr R. Motor adaptation as a process of reoptimization. J Neurosci. 2008;28: 2883–91. doi:10.1523/JNEUROSCI.5359- 07.2008

41. Balasubramanian R, Howe RD, Member S. Task Performance is Prioritized Over Energy Reduction.: 1–9.

42. Zeni JA, Richards JG, Higginson JS. Two simple methods for determining gait events during treadmill and overground walking using kinematic data. Gait Posture. 2008;27: 710–714. doi:10.1016/j.gaitpost.2007.07.007

43. Roemmich RT, Long AW, Bastian AJ. Seeing the Errors You Feel Enhances Locomotor Performance but Not Learning. Curr Biol. Elsevier Ltd.; 2016;26: 2707–2716. doi:10.1016/j.cub.2016.08.012

44. Leech KA, Roemmich RT. Independent voluntary correction and savings in locomotor learning. J Exp Biol. 2018; jeb.181826. doi:10.1242/jeb.181826

45. Selinger JC, Donelan JM. Estimating instantaneous energetic cost during non-steady-state gait. J Appl Physiol. 2014;117: 1406–1415. doi:10.1152/japplphysiol.00445.2014

46. Nakagawa S, Schielzeth H. A general and simple method for obtaining R2 from generalized linear mixed-effects models. Methods Ecol Evol. 2013;4: 133–142. doi:10.1111/j.2041-210x.2012.00261.x

